# Mitochondrial Calcium Uniporter protects hippocampal CA2 neurons from excitotoxic injury

**DOI:** 10.1101/2025.05.27.656314

**Authors:** Aleksandra Skweres, Małgorzata Beręsewicz-Haller, Omar Basheer, Martyna Nalepa, Aleksandra Owczarek, Maria Kawalec, Joanna Gruszczynska-Biegala, Barbara Zabłocka, Michał Węgrzynowicz

## Abstract

**Background:** The hippocampal region CA2, unlike neighboring CA1, is exceptionally resistant to excitotoxicity, although the mechanisms behind this phenotype are unknown. Given the importance of mitochondrial calcium buffering, we investigated whether the Mitochondrial Calcium Uniporter (MCU), recently found to be enriched in CA2, contributes to this resistance.

**Methods:** We employed immunostaining techniques in rodent brain tissue and organotypic slice cultures to visualize MCU distribution across hippocampal regions under both resting and excitotoxic conditions. Subsequently, we pharmacologically modulated MCU in an organotypic model of hippocampal excitotoxicity to assess its contribution to regional resistance to NMDA.

**Results:** We found a strong spatial correlation between resistance to excitotoxic injury and MCU expression. Notably, NMDA exposure resulted in MCU upregulation in CA2, and pharmacological inhibition of MCU sensitized CA2 neurons to excitotoxic damage in a dose-dependent manner, while having minimal effect on the already vulnerable CA1 neurons, which express low MCU levels.

**Conclusions:** MCU is known to exacerbate NMDA-induced cell injury, although our data indicate that CA2 neurons possess unique mitochondrial calcium handling capabilities enabling MCU to support neuroprotection. Our study provides novel insight into mechanisms supporting CA2 resistance to excitotoxic death and emphasizes context-dependent roles of MCU in neuronal injury or survival.

## Background

Due to its small size and difficulties in clearly delineate its borders in non-labelled hippocampus, the CA2 region has often been omitted from hippocampal studies. Despite limited knowledge, it has long been known that CA2 pyramidal neurons, unlike those in adjacent CA1 and CA3 regions, exhibit unusually high synaptic stability, manifested by their resistance to induction of long-term potentiation (LTP) and depression (LTD), particularly at their proximal dendrites (Chevaleyre and Siegelbaum 2010). For years, the molecular basis of this phenotype, similarly to specific role of CA2 in brain function remained unclear. However, recent emerging interest in this region has revealed its molecular uniqueness (Farris et al. 2019) and its critical importance for social recognition memory, the ability to distinguish individuals of the same species (Hitti and Siegelbaum 2014; Radzicki et al. 2024). Additionally, several CA2-specific cellular mechanisms have recently been demonstrated to underlie the limited plasticity of this region’s neurons. These include inhibition of the H-Ras/ERK/MAP2K signaling cascade by the CA2-enriched protein, regulator of G-protein signaling 14 (RGS-14) (Lee et al. 2010), a distinct structure of the perineuronal net, that, unlike in CA1 or CA3, surrounds both dendrites and spines of CA2 neurons (Carstens et al. 2016) and attenuation of postsynaptic cytosolic Ca^2+^ transients (Simons et al. 2009), a crucial mechanism in LTP/LTD formation. Given mitochondria’s role in neuronal Ca^2+^ buffering (White and Reynolds 1997), the last mechanism may also involve the efficiency of mitochondrial Ca^2+^ uptake via the Mitochondrial Calcium Uniporter (MCU) a protein recently found to be highly enriched in CA2 neurons and implicated in their distinct electrophysiological properties (Farris et al. 2019; Pannoni et al. 2023, 2025)

In addition to limited synaptic plasticity, CA2 pyramidal neurons are uniquely resistant to injury and death in various neurological disorders, particularly those involving excitotoxicity. Numerous studies report a consistent pattern of hippocampal damage: CA1 is severely affected, CA3 moderately, and CA2 minimally or not affected at all. This has been observed in post-mortem brains of patients with stroke (Bartsch et al. 2015), temporal lobe epilepsy (Steve et al. 2014) or traumatic brain injury (Maxwell et al. 2003) as well as in corresponding animal models(Yang et al. 2000; Beręsewicz-Haller et al. 2021). The cellular basis of CA2 resistance remains unclear. While it has been proposed that the mechanisms underlying synaptic stability, such as specific Ca^2+^ dynamics, may also contribute to the protection against excitotoxicity (Leranth and Ribak 1991; Dudek et al. 2016), due to shared dependence on postsynaptic N-methyl-D-aspartate (NMDA) receptor (NMDAR) activity, this hypothesis has not been experimentally confirmed. To investigate this, we used hippocampal organotypic rat slice cultures to examine the role of MCU in CA2 resistance to excitotoxic damage.

## Methods

### Animals

The studies were performed using C57Bl/6J mice and Wistar rats. Animals were housed in the Animal Facility of the Mossakowski Medical Research Institute, Polish Academy of Sciences (MMRI PAS). All experiments were carried out in accordance with European (EU Directive, 2010/63/EU) and Polish (Act no. 266/January 15, 2015) regulations.

### Organotypic hippocampal slice cultures

#### Establishment and maintenance of the cultures

Cultures were prepared as described by Beręsewicz-Haller and colleagues (Beręsewicz-Haller et al. 2021) with minor modifications. Hippocampi were isolated from 7-day-old rats and cut into 400 μm slices using a tissue chopper (McIlwain Tissue Chopper). Slices with preserved structure (verified under Primo Vert light microscope (Zeiss)) were placed on Millicell-CM membrane inserts (Millipore) and cultured at 36^0^C, 5% CO_2_ in humidified air. The culture medium contained 50% Neurobasal, 25% horse serum, 20% Hanks’ Balanced Salt Solution, B27 supplement, 1 M 4-(2-Hydroxyethyl)-1-piperazineethanesulfonic acid, 0.5 mM glutamax (all Gibco), glucose (5 mg/ml; Sigma-Aldrich) and Antibiotic Antimycotic Solution (Sigma-Aldrich). Serum was gradually withdrawn between day in vitro (DIV) 3 and DIV7, after which cultures were maintained serum-free.

### Treatment

On DIV8, prior to the experiment, slices were incubated with 2 µM PI (propidium iodide; Sigma-Aldrich) for 30 min. to label and exclude any damaged slices from further processing. In the remaining slices, excitotoxicity was induced by a 3 h exposure to NMDA (Sigma-Aldrich) at 25 µmol/l (selective injury) or 100 µmol/l (global injury control). MCU inhibition was performed with MCU-i4 (Tocris) at 2, 10 and 50 µmol/l for 4 h – either alone or with 25 µM NMDA. For the combined treatment, cultures were preincubated with MCU-i4 for 1 h, then co-incubated with NMDA for 3 h. Vehicle-treated slices served as negative controls. After treatment, NMDA- and MCU-i4-free medium was applied and slices were maintained for another 24 h. Following the treatment, slices were stained with 6 µM PI for 1 h to estimate neuronal damage under each condition.

### Immunohistochemistry

#### Immunostaining of fixed brain sections

Adult mouse and rat brain sections were prepared and stained as described by Nalepa et al. with slight modifications (Nalepa et al. 2025). Animals were perfused with 4% paraformaldehyde (Lach-Ner), and their brains, following cryoprotection, were sectioned at 30 μm, using a Hyrax M25 microtome (Zeiss) with an MTR freezing unit (Slee Medical).

Free-floating sections underwent antigen retrieval in 10 mM sodium citrate (Sigma-Aldrich), pH 6.0, at 80^°^C, blocking with 5% normal horse serum (Vector Laboratories) and quenching of endogenous peroxidases with 3% H_2_O_2_ (Chempur) in 20% methanol (Pol-Aura). Sections were then incubated with rabbit anti-MCU antibody (Cell Signaling Technology Cat# 14997, 1:1000), followed by biotinylated anti-rabbit secondary antibody (Vector Laboratories, 1:2000), ABC-HRP kit (Vector Laboratories), biotinylated tyramide (IRIS Biotech) and re-incubated with ABC-HRP. Signal was visualized using the HRP DAB Substrate Kit (Vector Laboratories). Sections were counterstained with cresyl violet (Acros Organics), mounted with DPX (Merck) and examined under an Axiovert.A1 microscope (Zeiss).

#### Staining of organotypic slices

Slices were fixed in methanol for 24 h at -20^°^C, then cut from the inserts together with underlaying membrane and stained as free-floating sections. After blocking, slices were incubated for 1 h at 37^°^C with either anti-MCU (1:500) or anti-RGS-14 (BioLegend Cat# 821801, 1:200). Next, sections were incubated with Alexa Fluor-conjugated secondary antibodies (488 nm anti-mouse or 647 nm anti-rabbit; Invitrogen, 1:1000). Nuclei were stained with DAPI (10 min.) and slices were mounted with FluorSave Reagent (Millipore).

### Confocal microscopy analysis

Microscopy was performed at the Laboratory of Advanced Microscopy Techniques, MMRI PAS. PI labelling was analysed at 514 nm using a confocal microscope (Zeiss LSM 510 Confocal Microscope), before and after slice exposure to NMDA and/or MCU-i4. Fixed organotypic slices stained for MCU or RGS-14 were analysed with a confocal microscope (Zeiss LSM 780 Confocal Laser Scanning Microscope) using 20x magnification. Images were processed with ZEN 3.5 Lite software. To assess protein (immunostaining) or damage (PI staining) distribution across CA1-CA3 subregions, fluorescence intensity was measured along a manually drawn line over the pyramidal layer, and plotted as signal intensity vs. distance. Mean PI signal intensities were calculated for CA1 and CA2 to compare damage between treatment groups.

### Statistics

Each culture was treated as a biological replicate, with values averaged from technical replicates (individual slices per condition per culture). Biological replicates were used for statistical analysis. Statistical analysis of obtained data was performed using GraphPad Prism 10 (GraphPad Prism). Two-way repeated-measure analysis of variance (ANOVA) was applied to identify effects of treatment, region or treatment x region interaction. Both factors – region and treatment were treated as within-subject variables, as measurements were obtained from both CA1 and CA2 under all concentrations within each of seven analysed cultures. Tukey’s post-hoc test was used for individual comparisons between experimental groups. A linear regression analysis was performed to evaluate the dose-response relationship between inhibitor concentrations and mean cell damage. The goodness-of-fit was assessed using the coefficient of determination (R^2^) and the p-value for the slope. Data are presented as means of biological replicates ± SEM. p < 0.05 was considered statistically significant.

## Results and Discussion

Recent studies report heterogeneous MCU mRNA(Farris et al. 2019) and protein (Pannoni et al. 2023) distribution across CA subregions in mice, with high expression in CA2, moderate in CA3 and low in CA1. We confirm this pattern in both mouse (Fig. 1A) and rat (Fig. 1B) brains, showing that MCU distribution correlates with pyramidal neurons resistance to pathology (see Introduction), suggesting a neuroprotective role of MCU.

**Fig 1.**
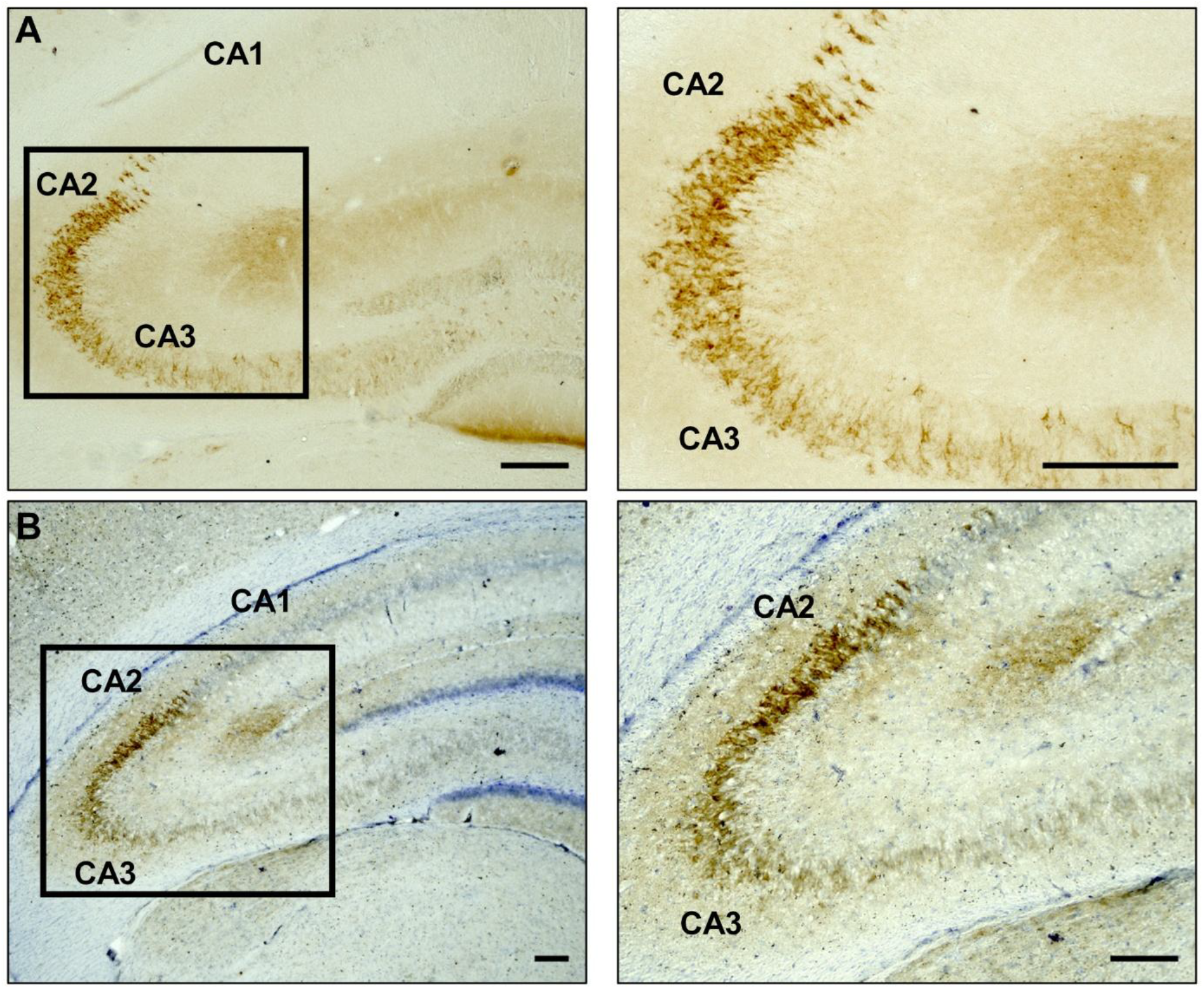
MCU protein distribution in mouse and rat hippocampus. **(A)** MCU immunostaining was performed on coronal sections of mouse hippocampus and visualized using diaminobenzidine (DAB, brown). Left: representative image showing CA1, CA2 and CA3 regions. Right: magnified view of the MCU-enriched area marked in the left panel. Scale bars: 200 µm. **(B)** MCU immunostaining was performed on coronal sections of rat hippocampus and visualized using DAB (brown). Nuclei were counterstained with cresyl violet (purple). Left: representative image showing CA1, CA2 and CA3 regions. Right: magnified view of the MCU-enriched area marked in the left panel. Scale bars: 200 µm.

For functional studies, we used a rat organotypic hippocampal slice culture model of excitotoxicity. Unlike dissociated cultures, organotypic slices preserve, to a large degree, hippocampal architecture and cell interactions (Stoppini et al. 1991). We first validated that our model retains region-specific sensitivity to excitotoxicity. Exposure to 25 μM NMDA caused strong increase in PI signal in pyramidal layer of CA1, indicating extensive neuronal damage, while CA2 neurons in the same slices remained largely unaffected. Further staining of the slices for RGS-14, a CA2 marker (Lee et al. 2010), confirmed the identity of the resistant region (Fig. 2A). To precisely visualize the distribution of injured neuron, we quantified PI fluorescence along pyramidal layer. PI intensity was high in CA1 and moderately high in CA3, while CA2 showed only baseline signal with no major peaks indicating severely damaged neurons (Fig. 2B). These findings demonstrate that our model reproduces the spatial pattern of excitotoxic damage seen in human post-mortem tissue and in vivo disease models.

**Fig 2.**
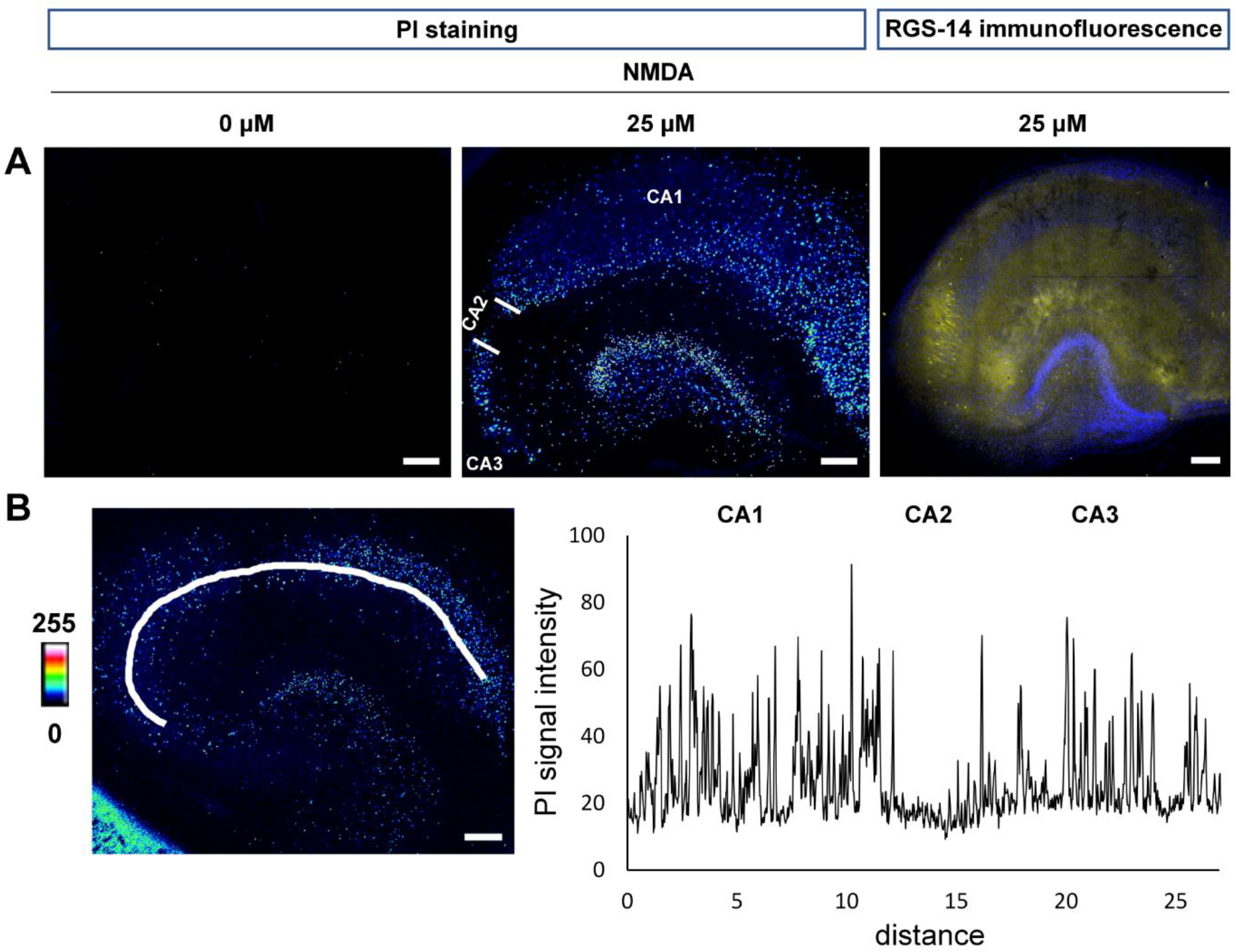
Differential sensitivity of hippocampal regions to excitotoxic stimuli in organotypic cultures. **(A)** Organotypic rat hippocampal slices were exposed to either vehicle (left) or 25 µM NMDA (middle and right) and stained with PI (pseudocolored for intensity) to visualize injured cells (left and middle). Following PI imaging, NMDA-treated slices were fixed and immunostained for the CA2 neuronal marker, RGS-14 (yellow) and stained with nuclei marker, DAPI (blue) (right). Representative images are shown. Scale bars: 200 µm. **(B)** Fluorescence intensity of PI staining was measured along a manually drawn line over the pyramidal layer in NMDA-treated slices (left; pseudocolored for intensity) and plotted as a function of distance (right). Representative image and corresponding graph are shown. Scale bar: 200 µm.

To assess whether MCU distribution in organotypic slices reflects its native pattern in adult rodent brains, we examined MCU localization in vehicle-treated slices. We found that its distribution along the pyramidal layer closely mirrored the in vivo pattern: CA1 neurons showed minimal MCU immunoreactivity, while CA2 expressed very high levels of this protein. CA3 neurons showed a slightly lower MCU signal, with intensity gradually decreasing with the distance from the CA2/CA3 border (Fig. 3A, 3B). These data show that, in addition to the spatial pattern of excitotoxic damage, MCU distribution is also preserved in organotypic cultures. Moreover, the strong correlation between regional MCU content and resistance to NMDA-induced injury supports a potential role for MCU in mediating region-specific neuroprotection.

**Fig 3.**
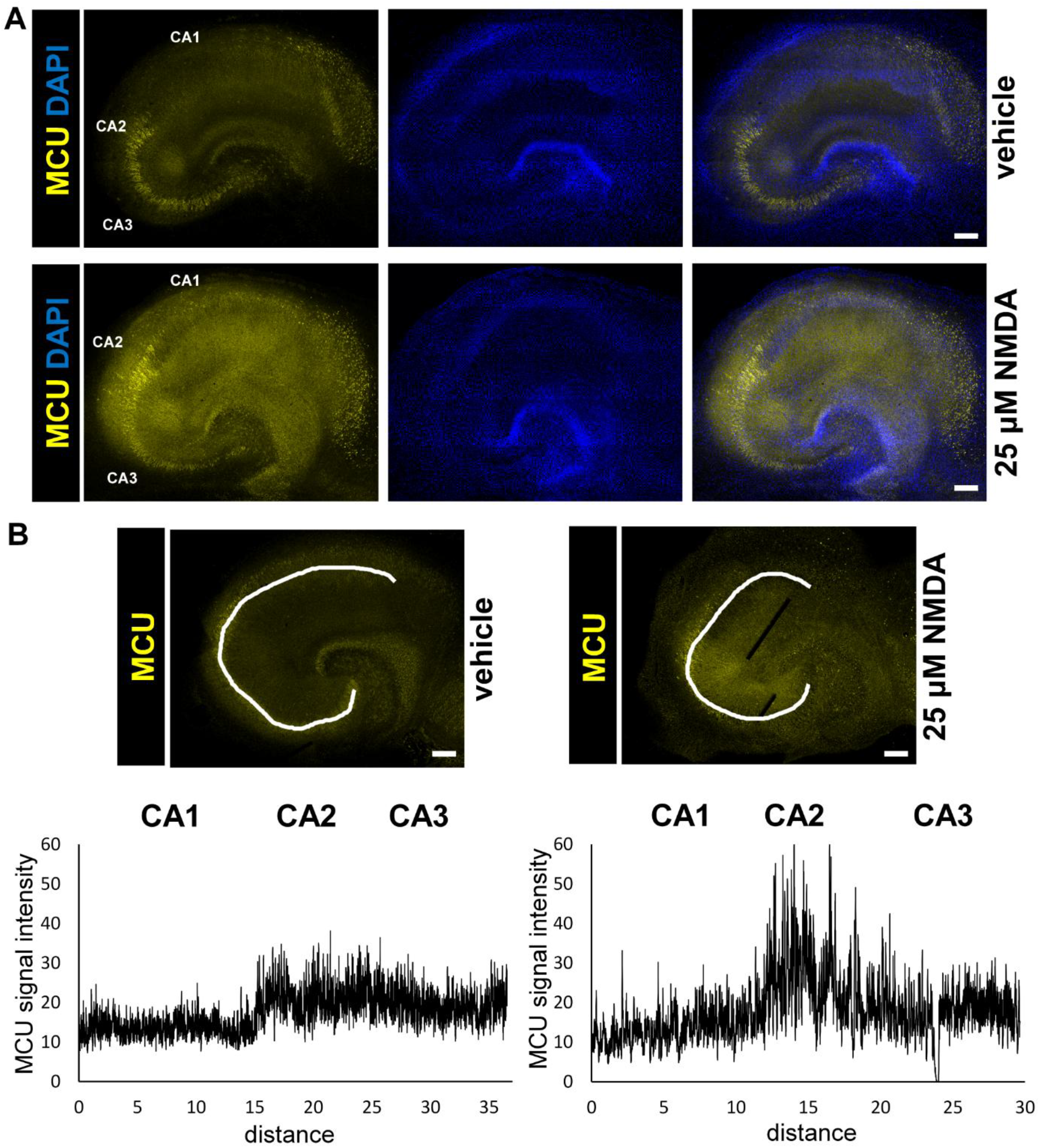
Changes in hippocampal MCU distribution in response to excitotoxic stimuli in organotypic cultures. **(A)** Organotypic rat hippocampal slices were exposed to either vehicle (top) or 25 µM NMDA (bottom) and, following fixation, immunostained for MCU (yellow, left), labelled with the nuclei marker, DAPI (blue, middle) and merged (right). Representative images are shown. Scale bars: 200 µm. **(B)** Intensity of MCU immunofluorescent labelling was measured along a manually drawn line over the pyramidal layer in vehicle-treated (left) and NMDA-treated (right) slices and plotted as a function of distance (bottom). Representative images and corresponding graphs are shown. Scale bars: 200 µm.

In slices exposed to 25 μM NMDA, we observed a marked upregulation of MCU specifically in CA2 neurons (Fig. 3A, 3B). Notably, this contrasts with previous findings in other neuronal populations, where MCU was shown to be downregulated in response to NMDA overstimulation as part of a protective feedback mechanism. For example, prolonged synaptic activity supresses MCU expression in excitotoxicity-prone cortical and hippocampal primary neurons thereby limiting mitochondrial Ca^2+^ overload and reducing neuronal damage (Qiu et al. 2013). The CA2-specific NMDA-induced increase in MCU expression suggests a distinct adaptive response in this region neurons, where MCU protein contributes to neuroprotection.

To test the hypothesis that MCU supports CA2 neuron survival, we analysed NMDA-induced injury in CA regions following inhibition of MCU using MCU-i4. This compound binds to the Mitochondrial Calcium Uptake 1 protein – an endogenous inhibitor of MCU that activates the channel in response to elevated extramitochondrial Ca^2+^ – and prevents its dissociation, thereby blocking MCU permeability even under high Ca^2+^ concentrations (Di Marco et al. 2020). First, we confirmed that MCU-i4 alone (2, 10 or 50 μM) caused no neuronal damage in CA1, CA2 or CA3 (Fig. 4A). However, when combined with 25 μM NMDA (a concentration non-damaging to CA2), MCU-i4 sensitized CA2 in a dose-dependent manner (Fig. 4A). Importantly, it appeared to have a minimal effect on CA1 damage induced by 25 μM NMDA, aligning with the low MCU expression in that region and confirming its selective action in MCU-enriched CA2 (Fig. 4A). Two-way repeated-measure ANOVA of PI intensity apart from treatment effect identified also an effect of region and treatment x region interaction, proving the distinct response of CA1 and CA2 neurons to MCU-i4 + NMDA exposure (Fig. 4B). Post hoc direct comparisons showed that PI intensity was consistently higher in CA1 than CA2 in slices exposed to 25 μM NMDA alone or with 2, 10 or 50 μM MCU-i4 despite MCU-i4 significantly increased PI signal in CA2 at 2 and 50 μM concentrations compared to NMDA alone (Fig. 4B). Notably, 2 μM MCU-i4 enhanced NMDA-induced damage in CA1, but this effect was absent at higher MCU-i4 concentrations (Fig. 4B).

**Fig 4.**
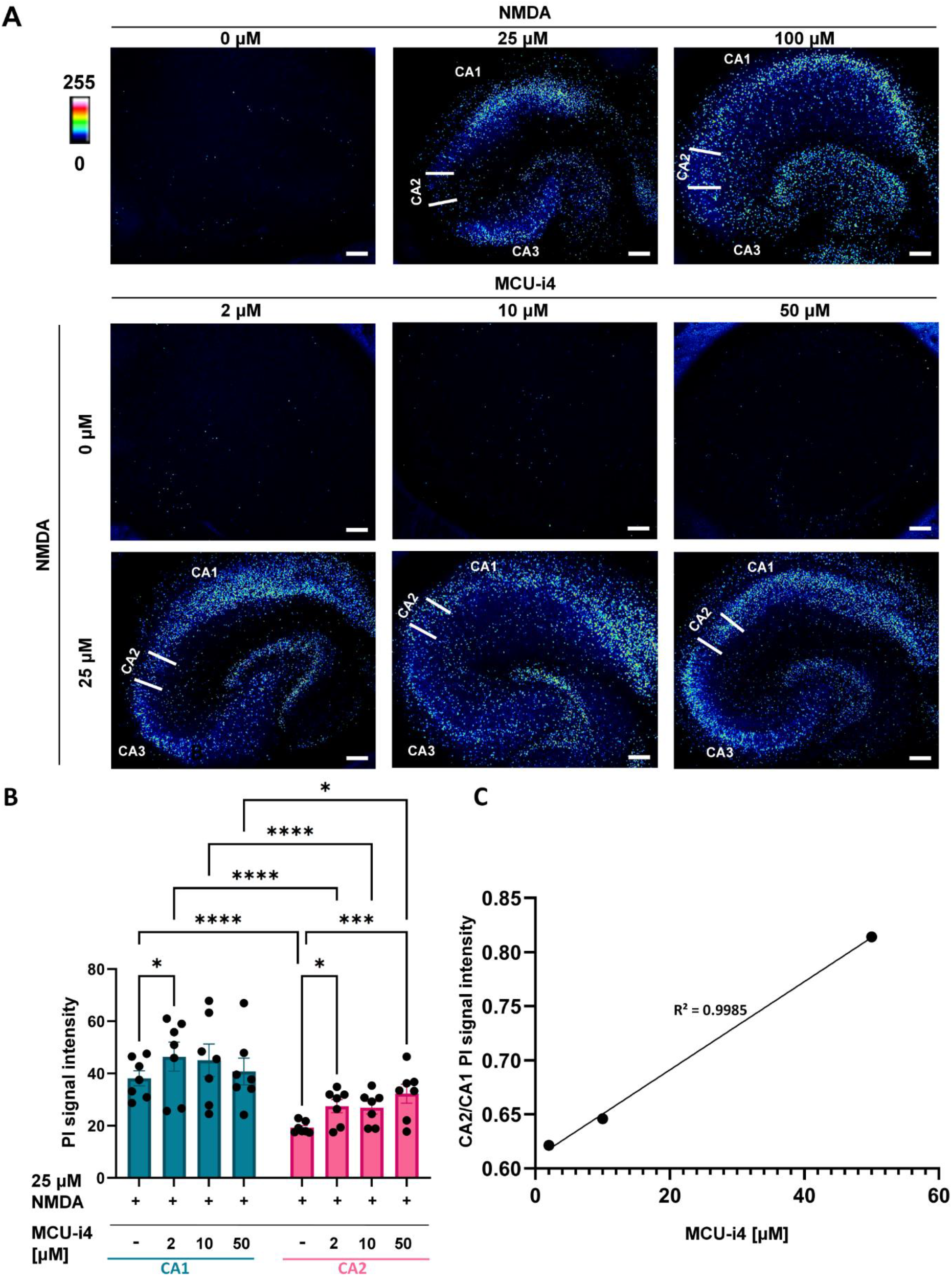
Effect of MCU inhibition on NMDA-induced damage in CA1 and CA2 regions in organotypic cultures. **(A)** Slices were exposed to vehicle (top left), NMDA (25 μM or 100 μM; top middle and right), MCU-i4 (2 μM; 10 μM or 50 μM; middle), or 25 μM NMDA combined with MCU-i4 (2 μM, 10 μM or 50 μM; bottom), and stained with PI (pseudocolored for intensity). Representative images are shown. Scale bars: 200 µm. **(B)** Average PI fluorescence intensity was quantified for CA1 and CA2 in slices exposed to 25 μM NMDA alone or in combination with MCU-i4 (2 μM, 10 μM or 50 μM). Two-way repeated-measure ANOVA revealed a main effect of treatment (F(3,18)=3,761; p=0.029), region (F(1,6)=40,69; p=0.0007) and a treatment x region interaction (F(3,18)=4.65; p=0.0141). Tukey’s test identified differences between CA1-25 μM NMDA vs CA1-25 μM NMDA + 2 μM MCU-i4 (p=0.0432), CA1-25 μM NMDA vs CA2-25 μM NMDA (p<0.0001), CA1-25 μM NMDA + 2 μM MCU-i4 vs CA2-25 μM NMDA + 2 μM MCU-i4 (p<0.0001), CA1-25 μM NMDA + 10 μM MCU-i4 vs CA2-25 μM NMDA + 10 μM MCU-i4 (p<0.001), CA1-25 μM NMDA + 50 μM MCU-i4 vs CA2-25 μM NMDA + 50 μM MCU-i4 (p=0.0371), CA2-25 μM NMDA vs CA2-25 μM NMDA + 2 μM MCU-i4 (p=0.0436) and CA2-25 μM NMDA vs CA2-25 μM NMDA + 50 μM MCU-i4 (p=0.0007). **(C)** Dose-response relationship was analyzed using the CA2/CA1 PI fluorescence intensity ratio to evaluate CA2 susceptibility relative to CA1 under increasing MCU-i4 concentrations combined with 25□μM NMDA. Mean CA2/CA1 ratios were 0.62□ ±□ 0.04, 0.65□ ±□ 0.05, and 0.81□ ±□ 0.06 for 25 µM NMDA + MCU-i4 at 2 µM, 10 µM, and 50□ µM concentrations, respectively. Linear regression revealed a positive correlation (R^2^□ =□ 0.9985, p□ =□ 0.025), indicating a linear dose-response relationship within the tested range. Data are presented as means ± SEM. n=7 biological replicates (individual cultures). *p < 0.05, ***p < 0.001, ****p < 0.0001.

To better control for culture-to-culture and imaging-to-imaging variability, we analysed the CA2/CA1 PI signal ratio as an internal reference across MCU-i4 doses combined with 25□ μM NMDA. Linear regression analysis revealed a significant positive correlation between inhibitor dose and the CA2-to-CA1 neuronal damage ratio (R^2^□ =□ 0.9985), indicating a strong linear dose-response relationship within the tested range and confirming a differential MCU-i4-mediated sensitization effect in CA2 compared to CA1 (Fig. 4C).

Our data contrast clearly with the common view that MCU activity via excessive entry of Ca^2+^ to mitochondria during excitotoxic events promotes neuronal injury and death. Prior studies with the use of different neuronal populations (Budd and Nicholls 1996; Stout et al. 1998) including hippocampal neurons (Qiu et al. 2013) showed that MCU knockout or inhibition reduced the excitotoxic damage, while its overexpression exacerbated cell injury. Our data indicate that this paradigm doesn’t apply universally, and in specific cell types, MCU may serve a protective function. We show that this applies specifically to the pyramidal neurons of CA2 region, and we don’t observe this effect in CA1 neurons, a population highly sensitive to excitotoxic insult. The discrepancy between our findings and earlier work may be due to the fact, that in the dissociated hippocampal neuronal cultures, CA2 neurons possibly lose their regional identity or, even if they preserve it, it is masked by overwhelming majority of neurons from other populations. Meantime, organotypic cultures preserve hippocampal subfield architecture, enabling detection of unique, region-specific neuroprotective mechanisms such as that of MCU in CA2.

It may be hypothesized that CA2 pyramidal neurons possess unique adaptations that enable them to tolerate high mitochondrial Ca^2^□ loads without triggering cellular dysfunction. Notably, CA2 neurons contain more mitochondria than CA1 neurons, both in the soma (Farris et al. 2019) and dendrites, where they are also larger in size than those in CA1 dendrites (Pannoni et al. 2023). This suggests greater mitochondrial Ca^2+^ buffering capacity. Additionally, compared to CA1, CA2 neurons exhibit enhanced cytosolic Ca^2^□ handling, including more efficient Ca^2+^ extrusion (Simons et al. 2009) improved buffering, and significantly higher expression of Ca^2^□ -binding proteins such as N-Terminal EF-Hand Calcium Binding Protein 1 (Sugita et al. 2002), calretinin (Seress et al. 1993), as well as parvalbumin and calbindin (Sloviter et al. 1991). These features may act jointly to reduce cytosolic Ca^2+^, even during prolonged cellular Ca^2+^ influx, such as during NMDAR overstimulation, thereby limiting extramitochondrial Ca^2+^ concentrations compared to CA1, thus protecting CA2 mitochondria from overload.

Why does inhibition of mitochondrial Ca^2^□ uptake kill CA2 neurons, while protecting other neuronal populations? This may be due to cytosolic Ca^2+^ overload. Given efficient Ca^2^□ extrusion, buffering and binding, CA2 neurons could be especially sensitive to free cytosolic Ca^2^□ accumulation. Cytosolic Ca^2+^ alone, independent of mitochondrial signalling, can trigger neurotoxic cascades via activation of Ca^2+^-dependent enzymes such as endonucleases or calpains (Orrenius et al. 1989; Gerencser et al. 2009). Thus, the differing responses of CA1 and CA2 neurons to excitotoxicity and MCU inhibition may stem from varying sensitivities to cytosolic versus mitochondrial Ca^2+^ accumulation.

Although limited, some studies report neuroprotective roles of MCU. For instance, MCU protected phenochromocytoma PC12 cells from 1-methyl-4-phenylpyridinium-induced death (Wang et al. 2018). Also, glucocerebrosidase 1 knockout led to reduced MCU expression and greater glutamate-induced cytosolic Ca^2+^ accumulation, exacerbating cell damage (Plotegher et al. 2019). Notably, both models relate to Parkinson’s disease, suggesting a potential role of MCU loss in its pathogenesis. These findings along with ours, indicate that the role of MCU in neuronal damage is not as unambiguous as it is commonly believed, and varies by cell type and condition. Therefore, any therapeutic strategies targeting MCU should consider the potential harmful effects on specific cell populations.

## Conclusions

Our findings identify a novel mechanism behind CA2 resistance to excitotoxic stimuli, that involves MCU. We show that MCU levels are inversely correlated with excitotoxic sensitivity across hippocampal CA subregions, with CA2 showing the highest MCU expression and lowest vulnerability. MCU levels in CA2 further increases after NMDA exposure, suggesting stress-induced upregulation and potential involvement in CA2 neuron protection. Supporting this, MCU inhibition suppresses CA2 resistance, making it vulnerable to NMDA. These results establish MCU as a CA2-specific neuroprotective factor involved in the long-known resistance of CA2 neurons to excitotoxicity (Dudek et al. 2016).

## List of abbreviations

ANOVA: analysis of variance
DAB: diaminobenzidine
DIV: day in vitro
LTD: long-term depression
LTP: long-term potentiation
MCU: mitochondrial calcium uniporter
NMDA: N-methyl-D-aspartate
NMDAR: N-methyl-D-aspartate receptor
PI: propidium iodide
RGS-14: Regulator of G-protein signaling 14

## Declarations

### Ethics approval and consent to participate

All experiments were carried out in accordance with European (EU Directive, 2010/63/EU) and Polish (Act no. 266/January 15, 2015) regulations.

### Consent for publication

Not applicable

### Availability of data and materials

The datasets generated and analysed during the current study are available in the RepOD repository, https://repod.icm.edu.pl/dataset.xhtml?persistentId=doi:10.18150/P9YKB2.

## Author’s contribution

**AS** acquired funding; administered the project; developed the methodology; performed the experiments; supervised the experimenters; analysed, validated and visualized the data; wrote the original draft of the manuscript; **MBH** provided the methodology; performed the experiments; edited the manuscript; **OB** performed the experiments; **MN** performed the experiments; **AO** performed the experiments; **MK** performed the experiments; **JGB** acquired funding; performed the experiments; edited the manuscript; **BZ** conceived the project; provided the methodology and resources; supervised the experimenters; edited the manuscript; **MW** acquired funding; conceived and administered the project, provided methodology and resources; supervised the experimenters and analysts; analysed, validated and visualized the data; wrote the original draft of the manuscript. All authors read and approved the final manuscript.

## Acknowledgements

Not applicable

